# Molecular and functional characterization of telomeric repeat-containing RNAs in Chinese hamster ovary cells

**DOI:** 10.64898/2026.04.01.715793

**Authors:** Beatriz Domingues-Silva, Claus M. Azzalin

## Abstract

Mammalian telomeric DNA comprises long tracts of tandem TTAGGG repeats. The same repeats are also found at internal chromosomal regions called interstitial telomeric sequences (ITSs). Telomeres are transcribed into UUAGGG-containing transcripts, named TERRA, which serve multiple functions in maintaining telomere integrity. Complementary RNAs containing C-rich telomeric repeats, named ARIA, have also been identified in few yeast mutants and mammalian cells with dysfunctional telomeres. The molecular features and functions of ARIA remain understudied, mainly due to its low abundance and the lack of suitable cellular systems. Here, we show that Chinese hamster ovary (CHO) cells produce abundant TERRA and ARIA transcripts, predominantly originating from ITSs. Both RNAs are polyadenylated, exhibit relatively short half-lives and form large cellular foci. We also show that ARIA depletion leads to exposure of single-stranded (ss) DNA at ITSs and that ssDNA exposure increases when ITS DNA is damaged. SsDNA formation does not require the DNA damage signaling kinases ATM and ATR, nor the exonucleases DNA2 and EXO1; however, ATM prevents excessive ssDNA accumulation when ARIA function is inhibited. These findings establish CHO cells as a powerful model to dissect telomeric RNA functions and reveal ARIA as a key regulator of telomeric repeat DNA integrity.

## Introduction

Telomeres are located at the physical ends of linear eukaryotic chromosomes. In mammals, they comprise long tracts of the double stranded (ds) 5ʹ-TTAGGG-3ʹ/5ʹ-CCCTAA-3ʹ DNA sequence, with the G-rich strand protruding over its complement to form the so-called G-overhang (1,2). Bound to the multi-protein complex shelterin, telomeres safeguard genome stability by preventing spurious activation of DNA damage repair machineries at chromosome ends (1). Moreover, in cells lacking mechanisms of de novo synthesis of telomeric DNA, such as human primary cells, telomeres set the cell lifespan by shortening at each cell division until they trigger replicative senescence or death (3).

Stretches of telomeric repeats are also found at internal chromosomal regions, referred to as interstitial telomeric sequences (ITSs) (4–6). Cytogenetic and bioinformatics analyses classify ITSs in vertebrates into two main groups: short ITSs and heterochromatic ITSs (5,6). Short ITSs consist of arrays of TTAGGG repeats covering up to a few hundred base pairs and are interspersed at various chromosomal positions. In the human genome, thousands of s-ITSs are found across all chromosomes and usually contain 2–25 repeat units (7,8). Short ITSs are thought to arise from the insertion of telomeric repeats during DNA double-strand break (DSB) repair, via mechanisms involving non-homologous end joining and the activity of the reverse transcriptase telomerase (8–10). Heterochromatic ITSs comprise large blocks of telomeric repeats, often located in centromeric or pericentromeric regions and spanning hundreds of kilobases. They are proposed to originate from chromosome fusion events during karyotype evolution, followed by rearrangements including breakage and amplification (5). While the human genome lacks conventional heterochromatic ITSs, they have been described in several mammals, amphibians, reptiles, fishes and birds (5,8). Well-documented heterochromatic ITSs are present in the genome of the Chinese hamster (*Cricetulus griseus*), where they comprise extended arrays of both conserved and degenerated TTAGGG repeats spanning 250–500 kb across several chromosomes and accounting for approximately 5% of the genome (11–16). ITSs are preserved in immortalized Chinese hamster cell lines, including Chinese hamster ovary (CHO) cells. In these cells, telomeres are estimated to be shorter than 1 kb; hence, the vast majority of TTAGGG repeats in CHO cells originate from intrachromosomal regions (16).

Despite being heterochromatic, telomeres are transcribed into the RNA TERRA, which is conserved across virtually all eukaryotes, including humans, rodents, fish, birds, nematodes, yeasts and plants (17). In humans, RNA polymerase II initiates TERRA transcription from CpG-rich subtelomeric promoters towards chromosome ends, generating transcripts that include chromosome-specific subtelomeric sequences followed by multiple UUAGGG repeats (18–22). Human TERRA carries a 5ʹ 7-methylguanosine cap and is partly 3ʹ-end polyadenylated. In HeLa cancer cells, roughly 10% of TERRA is poly(A)+, is found mainly soluble in the nucleoplasm and has a half-life of more than 8 hours. In contrast, poly(A)- TERRA is largely associated with telomeric chromatin through interactions with shelterin proteins and the formation of DNA:RNA hybrids (telomeric R-loops), and exhibits a substantially shorter half-life (21,23–27). Functionally, TERRA is central to various nuclear pathways that maintain telomere integrity, such as facilitating telomeric DNA replication, promoting heterochromatin deposition at telomeres and regulating telomere elongation via the activity of telomerase or homology-directed repair (27–34). TERRA has also been shown to stimulate the activation of DNA repair activities at dysfunctional telomeres (35).

Beyond TERRA, other transcripts produced from chromosome ends have been identified, including ARIA, a C-rich RNA made of repeats complementary to TERRA repeat sequence. ARIA has been mainly characterized in fission yeast mutant strains lacking the shelterin proteins Taz1p and Rap1p, and its conservation and functions across eukaryotes remain poorly understood (17,36,37). In mammalian cells, ARIA is barely detected under normal conditions (18,21). However, in cells with dysfunctional telomeres, damage-induced long noncoding RNAs (dilncRNAs) containing CCCUAA and UUAGGG repeats are produced from telomeres by RNA polymerase II to mediate DNA damage signaling (38). Like telomeres, ITSs can also be transcribed. In humans, many telomeric repeat-containing transcripts originate from short ITSs, which are transcribed in either direction, producing G-rich TERRA-like and C-rich ARIA-like RNAs (22,39).

The initial study that reported the discovery of TERRA found that CHO cells display exceptionally high TERRA levels, much higher than in human cells (18). However, this observation has not been investigated further. Here we show that CHO cells produce highly abundant TERRA and ARIA transcripts, likely derived mainly from large ITSs. We also provide the first molecular characterization of both CHO TERRA and ARIA and demonstrate that ARIA supports the integrity of ITS DNA. Our findings establish CHO cells as a novel and uniquely suited model system to study mammalian telomeric repeat-containing RNAs.

## Materials and Methods

### Cell culture, manipulation and analysis

CHO cells (kind gift from Elena Giulotto, University of Pavia, Italy) were cultured in high glucose DMEM, GlutaMAX (Thermo Fisher Scientific), supplemented with 5% tetracycline-free fetal bovine serum (Pan BioTech), 100 U/ml penicillin-streptomycin (Thermo Fisher Scientific) and non-essential amino acids (Thermo Fisher Scientific). Cells were confirmed to be negative for mycoplasma contamination using the Lookout® Mycoplasma PCR Detection Kit (Sigma-Aldrich). When indicated, cells were treated with 5 µg/ml actinomycin D (Sigma-Aldrich), 150 nM or 1 µM camptothecin (Selleckchem), 10 µM KU-55933 (ATM inhibitor, Selleckchem), or 10 µM VE-821 (ATR inhibitor, Selleckchem). Antisense LNA GapmeRs (ASOs; Qiagen) and DsiRNAs (Integrated DNA Technologies) were transfected into cells using the Lipofectamine RNAiMAX reagent (Invitrogen) at a final concentration of 20 nM. ASO sequences were: 5ʹ-ACCTACCTCAATACCA-3ʹ for ASO Ctrl (LG00787372-DDA); 5ʹ-TAACCCTAACCCTAAC-3ʹ for ASO α-TERRA (LG00195219-DDA); 5ʹ-TTAGGGTTAGGGTTA-3ʹ for ASO α-ARIA (LG00196336-DDA). DsiRNA target sequences were: 5ʹ-CAGUCAUCUGAUAGUUGCAA-3ʹ for siDNA2-2; 5ʹ-CAGUUGAAGAACAGGUGGAU-3ʹ for siDNA2-3; 5ʹ-CUGGAAUGACAAAUCAUGUC-3ʹ for siEXO1-2; 5ʹ-AGAAAGCUAACAAGAAGAAU-3ʹ for siEXO1-3. The siCtrl was a non-targeting DsiRNA (51-01-14-04). For TRF1-FokI expression, a TRF1-FokI cDNA (kind gift from Roger Greenberg, University of Pennsylvania, Philadelphia, USA) was cloned into the lentiviral vector pLVX-TetOne-Puro (Clontech), followed by in frame insertion of a gBlock containing a Flag tag and an eGFP cDNA (Integrated DNA Technologies) at the N-terminus of TRF1-FokI. A gBlock containing only eGFP was also cloned into pLVX-TetOne-Puro to generate a control plasmid. The obtained plasmids, pLVX-Flag-GFP-TRF1-Fok1 and pLVX-eGFP, were used to produce lentiviruses and infect CHO cells according to standard procedures. Infected cells were selected 24 h after infection in medium supplemented with 10 µg/ml puromycin (Sigma-Aldrich). Ectopic proteins expression was induced with 200 ng/ml doxycycline (Sigma-Aldrich) for 24 h. For cell proliferation analysis, cells were washed with 1× PBS, trypsinized and counted at regular intervals using a hemocytometer. For viability assays, total cells (adherent and floating in the medium) were collected, washed twice with 1× PBS, and incubated in 10 µg/ml propidium iodide (Sigma-Aldrich) in 1× PBS at 4 °C for 10 min. For cell cycle analysis, cells were washed once with 1× PBS, trypsinized and fixed in ice-cold 70% ethanol for 20 min. Cells were then treated with 25 µg/ml RNase A (NZYTech) in 1× PBS at 37 °C for 20 min and stained with propidium iodide as above. Flow cytometry was performed at the Flow Cytometry platform of GIMM using a BD Accuri C6 instrument (BD Biosciences). Data were analyzed using FlowJo software.

### RNA preparation and analysis

Total RNA was isolated using TRIzol reagent (Invitrogen) following standard procedures, treated twice with 13.6 Kunitz units of DNase I (Qiagen), resuspended in RNase-free water and stored at −80 °C. For polyadenylated RNA purification, 10-20 µg of RNA were diluted in 100 µl of RNase-free water and an equal volume of binding buffer (20 mM Tris-HCl, pH 7.5, 1 M LiCl, 2 mM EDTA) was added. Samples were heated to 75 °C for 4 min, placed on ice for 2 min and incubated with oligo(dT)25 Magnetic Beads (New England Biolabs) at room temperature for 1 h on a rotating wheel. The flow-through, corresponding to poly(A)- RNA, was precipitated with 1/10 volume of 3 M sodium acetate (pH 5.2) and 3 volumes of 100% ethanol at −20 °C overnight and resuspended in 20 µl of RNase-free water. Poly(A)+ RNA bound to the beads was washed three times for 5 min with wash buffer I (20 mM Tris-HCl pH 7.5, 0.5 M LiCl, and 2 mM EDTA) and once for 5 min with low-salt buffer (20 mM Tris-HCl pH 7.5, 0.2 M LiCl, and 2 mM EDTA). 20 µl of elution buffer (10 mM Tris-HCl pH 7.5) were added to the beads and samples were heated at 80 °C for 3 min. For northern blot analysis, 5-15 µg of RNA were separated on 1.2% agarose gels containing 0.7% formaldehyde and 150 ng/ml ethidium bromide (Sigma-Aldrich) in 1× FA buffer (20 mM MOPS, 5 mM NaOAc, 10 mM EDTA, pH 7), followed by blotting onto nylon membranes (Hybond N+, Cytiva) by capillary transfer and UV cross-linking using a UV Stratalinker 2400 (Stratagene). TERRA and ARIA hybridizations were carried out at 55 °C overnight with a double-stranded telomeric probe (Telo2 probe; (18)) radioactively labeled using the Klenow fragment (New England Biolabs) and [α-32P]dCTP to detect TERRA or [α-32P]dGTP (PerkinElmer) to detect ARIA. Post-hybridization washes were performed twice in 2× SSC (0.3 M NaCl, 0.03 M trisodium citrate, pH 7.2), 0.2% SDS for 20 min and once in 0.2× SSC, 0.2% SDS for 30 min at 55 °C. Radioactive images were acquired using an Amersham Typhoon IP imager and analyzed using ImageJ. To detect 18S rRNA, β-Actin and U6 (loading and fractionation controls), the same membranes were stripped with boiling 0.1% SDS and re-hybridized with oligonucleotides 5ʹ-end radiolabeled using T4 polynucleotide kinase (New England Biolabs) and [γ-32P]ATP (PerkinElmer) at 50-55 °C overnight. Oligonucleotide sequences were: 5ʹ-CCATCCAATCGGTAGTAGCG-3ʹ for 18S (Integrated DNA Technologies); 5ʹ-TGGTACGACCAGAGGCATACAG-3ʹ for β-Actin (Sigma-Aldrich); 5ʹ-GGAACACTTTACGAATTTGCGT-3ʹ for U6 (Integrated DNA Technologies). Post-hybridization washes were performed twice in 2× SSC, 0.2% SDS for 20 min and once in 1× SSC, 0.2% SDS for 30 min at 50-55 °C. Image acquisition and analysis were as above. For dot-blot analysis, 1-2 µg of RNA were diluted in 100 µl of RNase-free water and denatured at 65 °C for 7 min, followed by dot-blotting onto nylon membranes and UV cross-linking. Hybridizations were carried out at 55 °C overnight with 5ʹ-end radiolabeled telomeric oligonucleotides. Sequences were: 5ʹ-(CCCTAA)5-3ʹ for TERRA and 5ʹ-(TTAGGG)5-3ʹ for ARIA (Integrated DNA Technologies). Post-hybridization washes were performed twice in 2× SSC, 0.2% SDS for 20 min and once in 0.2× SSC, 0.2% SDS for 30 min at 55 °C. After image acquisition, membranes were stripped and re-hybridized with radiolabeled 18S and β-Actin oligonucleotides. Image acquisition and analysis were as above. For RT-qPCR analysis, 2 µg of total RNA were reverse transcribed with SuperScript II or IV reverse transcriptase (Invitrogen) using oligo(dT)30 (Integrated DNA Technologies). qPCR was performed using the iTaq Universal SYBR Green Supermix (Bio-Rad) and a 7500 Fast Real-Time PCR instrument (Applied Biosystems). Oligonucleotide sequences were: 5ʹ-TCCTTGGGGCGGTGGAAAAC-3ʹ, c-Myc Forward; 5ʹ-GGCTGCACCGAGTCGTAGTC-3ʹ, c-Myc Reverse (Sigma-Aldrich); 5ʹ-AGAGGGCATGCTGTCCAAAA-3ʹ, DNA2 Forward; 5ʹ-TTGGGAGCAGGCATTAACCA-3ʹ, DNA2 Reverse; 5ʹ-GGATAAGCGTTCCCTACTGGG-3ʹ, EXO1 Forward; 5ʹ-ATGGGGATTCGCCTCCAATG-3ʹ, EXO1 Reverse (Integrated DNA Technologies); 5ʹ-GCTCTCTGCTCCTCCCTGTTC-3ʹ, GAPDH Forward; 5ʹ-AACCAGGCGTCCAATACGGC-3ʹ, GAPDH Reverse (Sigma-Aldrich).

### DNA preparation and analysis

Genomic DNA was isolated by phenol:chloroform extraction following standard procedures. 5-15 µg of DNA were digested with HinfI and HindIII (New England Biolabs) at 37 °C overnight and separated on 0.7% agarose gels containing GelRed (Biotium) in 0.5× TBE buffer. Gels were dried and hybridized at 50 °C with radiolabeled Telo2 probe to detect single-stranded G-rich telomeric DNA and with a 5ʹ-end radiolabeled telomeric oligonucleotide to detect single-stranded C-rich telomeric DNA. Post-hybridization washes were performed twice in 2x SSC, 0.5% SDS for 20 min and once in 1x SSC, 0.5% SDS for 30 min at 50 °C. After radioactive image acquisition, gels were incubated in denaturation solution (1.5 M NaCl, 0.5 M NaOH) at room temperature for 30 min and in neutralization solution (0.5 M Tris-HCl pH 7.5, 3 M NaCl) for 15 min, followed by hybridization as above. Image acquisition and analysis were performed as for northern blotting.

### Protein preparation and analysis

Total proteins were extracted by lysing cells in 2× Laemmli buffer (4% SDS, 20% glycerol, 120 mM Tris-HCl pH 6.8), followed by homogenization through a 27G syringe needle and boiling at 95 °C for 5 min. 15-30 µg of proteins were mixed with SDS loading buffer (50 mM Tris-HCl pH 6.8, 2% SDS, 6% glycerol, 0.005% bromophenol blue and 1% β-mercaptoethanol), denatured at 95 °C for 5 min and separated on 10% or 14% polyacrylamide gels. Proteins were transferred onto nitrocellulose membranes (Amersham Protran 0.45 NC, Cytiva) using a Trans-Blot SD Semi-Dry Transfer Cell apparatus (Bio-Rad). Membranes were blocked at room temperature for 1 h in blocking solution: 5% milk in 1×PBS with 0.1% Tween-20 (PBS-T) for unmodified epitope-specific antibodies or 5% BSA in 1× PBS-T for phosphorylated epitope-specific antibodies. Membranes were then incubated with primary antibodies diluted in blocking solution at 4 °C overnight. After three washes of 10 min each in 1× PBS-T, membranes were incubated with HRP-conjugated secondary antibodies diluted in blocking solution at room temperature for 1 h and washed as before. Chemiluminescent signals were detected using ECL reagents (Cytiva) and an Amersham 680 RGB imaging system. The following primary antibodies were used: a mouse monoclonal anti-γ-H2AX (Millipore, 05-636, 1:2000), a mouse monoclonal anti-GFP (Roche, 11814460001, 1:1000), a rabbit monoclonal anti-pCHK1 (Cell Signaling Technology, 2348, 1:500), a mouse monoclonal anti-β-Actin (Santa Cruz Biotechnology, sc-47778, 1:2000), a rabbit monoclonal anti-Vinculin (Cell Signaling Technology, 13901, 1:1000). Secondary antibodies were HRP-conjugated goat anti-mouse and anti-rabbit IgGs (Novus Biologicals, NB7539 and NB7160, 1:3000).

### DNA fluorescence in situ hybridization (FISH) on metaphase spreads

Cells were treated with 200 ng/ml Colchicine (Sigma-Aldrich) for 4h and mitotic cells were harvested by shake-off and centrifuged at 400 x g at 10 °C for 4 min. Cell pellets were resuspended in 7 ml of pre-warmed hypotonic solution (0.075 M KCl), incubated at 37 °C for 10 min, followed by centrifugation and cell fixation in 7 ml of ice-cold fixative solution (methanol:acetic acid 3:1, Supelco) at room temperature for 20 min. After a second wash in fixative solution, cells were concentrated by centrifugation, spread on microscopy slides and air dried. Slides were incubated in 1x PBS containing 20 µg/ml RNase A at 37 °C for 30 min, rinsed in 1x PBS and fixed with 4% formaldehyde in 1x PBS at room temperature for 2 min. Slides were then incubated in pre-warmed 2 mM glycine pH 2 supplemented with 70 µg/ml pepsin (Sigma-Aldrich) at 37 °C for 5 min, rinsed in 1x PBS, fixed again with 4% formaldehyde in 1x PBS at room temperature for 2 min and dehydrated through an ethanol series (70%, 90% and 100%) for 5 min each. A TYE 563-conjugated G-rich telomeric LNA probe (5ʹ-TYE563-TTAGGGTTAGGGTTAGGG-3ʹ; Exiqon) diluted to 20 nM in hybridization solution (2x SSC, 50% formamide, 0.5% blocking solution (Roche)) was applied onto slides followed by denaturation on a heating plate at 80 °C for 3 min and hybridization at room temperature for 2 h in a dark humid chamber. Slides were washed twice in 2x SSC for 10 min, once in 2x SSC containing 100 ng/ml DAPI (Sigma-Aldrich) for 5 min and rinsed in 2x SSC at room temperature. After another dehydration through an ethanol series as above, slides were air-dried and mounted in vectashield (Vector Laboratories). Images were acquired at the Bioimaging Platform of GIMM with a widefield Zeiss Axio Observer equipped with a cooled CCD Axiocam 506 mono camera and using a 63x/1.4 NA oil Ph3 M27 Plan-Apochromat objective. Image analysis was performed using ImageJ.

### Native DNA FISH

Cells grown on coverslips were rinsed twice with ice-cold 1x PBS, fixed with 3.6% formaldehyde in 1x PBS at room temperature for 10 min, rinsed twice with 1x PBS, permeabilized with CSK buffer (100 mM NaCl, 300 mM sucrose, 3 mM MgCl2, 0.5% Triton X-100 and 10 mM PIPES pH 7) at room temperature for 7 min and rinsed twice with 1x PBS. Slides were then incubated in 20 µg/ml RNase A in 1x PBS at 37 °C for 45 min, rinsed three times with 1x PBS and once with 2x SSC. Cells were dehydrated through an ethanol series (70%, 95% and 100%) for 5 min each and coverslips were let to air-dry. Coverslips were placed on microscope slides with 20 nM TYE563-conjugated G-rich telomeric LNA probe in hybridization solution (2x SSC, 30% formamide, 0.5% blocking solution) and incubated at room temperature for 2 h in a dark humid chamber. Coverslips were washed twice in 2x SSC for 10 min, once in DAPI in 2x SSC for 5 min, rinsed in 2x SSC and mounted on microscope slides with vectashield. Image acquisition was performed as for DNA FISH on metaphase spreads. The numbers of foci were counted manually using ImageJ, intensities of the foci were quantified using a semiautomated ImageJ macro kindly provided by Emilio Cusanelli (University of Trento, Trento, Italy).

### Indirect immunofluorescence (IF) and IF combined with DNA FISH (IF/DNA FISH)

Cells grown on coverslips were rinsed twice with ice-cold 1× PBS and pre-extracted with CSK buffer on ice for 7 min, fixed with 3.6% formaldehyde in 1× PBS at room temperature for 10 min and re-permeabilized with CSK buffer at room temperature for 7 min. After rinsing twice with 1× PBS, cells were incubated in blocking solution (5 mg/mL BSA in 1× PBS-T) for 45 min at room temperature. Primary antibodies were diluted in blocking solution and incubated for 1 h at room temperature in a humid chamber, followed by three washes with 1× PBS-T for 5 min. Secondary antibodies were incubated for 40 min at room temperature, followed by three washes with 1x PBS-T for 5 min. DNA was stained with DAPI for 5 min and coverslips were mounted in vectashield. For IF/DNA FISH, cells were pre-extracted with CSK buffer, fixed with 2% formaldehyde, permeabilized with 0.5% Triton X-100 for 5 min and then fixed with ice-cold methanol at −20 °C for 20 min. Blocking solution was supplemented with 20 mM glycine and 20 µg/ml RNase A and incubation was performed at 37 °C for 30 min. After primary and secondary antibody incubations, an additional fixation in 4% formaldehyde was performed for 10 min before hybridization. Slides were incubated with 20 nM AF594-conjugated G-rich LNA probe (5ʹ-AF594-TTAGGGTTAGGGTTAGGG-3ʹ; Exiqon) in hybridization solution (2× SSC, 50% formamide, 0.5% blocking solution), denatured at 80 °C for 5 min and incubated for 2 h at room temperature in a dark humid chamber, followed by two washes in 2× SSC for 10 min. DNA was stained with DAPI and coverslips were mounted in vectashield. Primary antibodies were a rabbit polyclonal anti-γ-H2AX (Abcam, ab2893, 1:200), a rabbit polyclonal anti-Flag (Sigma-Aldrich, F7425, 1:500), a mouse monoclonal anti-Flag (Sigma-Aldrich, F1804, 1:500). Secondary antibodies were donkey anti-rabbit and donkey anti-mouse IgGs conjugated with Alexa Fluor 568 or 488 (Invitrogen, 1:1000). Image acquisition and analysis was performed as for DNA FISH on metaphase spreads.

### RNA FISH

Cells grown on coverslips were rinsed twice with ice-cold 1x PBS and pre-extracted with CSK buffer supplemented with 10 mM Vanadyl Ribonucleoside Complex (New England Biolabs) on ice for 7 min. Cells were rinsed twice with ice-cold 1x PBS, fixed with 4% formaldehyde in 1x PBS at room temperature for 10 min and rinsed twice with 1x PBS. Cells were re-permeabilized with CSK buffer at room temperature for 10 min and rinsed twice with 1x PBS, followed by dehydration through an ethanol series (70%, 90% and 100%) for 5 min each. For TERRA detection, a Cy3-conjugated C-rich telomeric PNA probe (5ʹ-Cy3-OO-CCCTAACCCTAACCCTAA-3ʹ; Panagene) diluted to 10 nM in TERRA hybridization solution (50% formamide, 10% dextran sulfate (Sigma-Aldrich), 2 mg/ml BSA, 2x SSC) was applied onto slides. For ARIA detection, a TYE 563-conjugated G-rich telomeric LNA probe diluted to 20 nM in ARIA hybridization solution (30% formamide, 10% dextran sulfate, 2 mg/mL BSA, 2x SSC) was used. Slides were covered with coverslips and incubated at room temperature for 2 h in a light protected humid chamber and then rinsed with 2x SSC. For TERRA detection, the washes were three times with 2x SSC/50% formamide at 39 °C for 5 min, three times with 2x SSC at 39 °C for 5 min and once with 2x SSC at room temperature for 5 min. For ARIA detection, the washes were two times with 2x SSC at room temperature for 10 min. DNA was stained with DAPI and coverslips were mounted on microscopy slides in vectashield. Images were acquired at the Bioimaging Platform of GIMM with a confocal point-scanning Zeiss LSM 710 or Zeiss LSM 880 using a 63x/1.4 NA oil DIC M27 Plan-Apochromat objective. Image analysis was performed using ImageJ.

### Statistical analysis

Statistical analysis was performed in GraphPad Prism 8. For direct comparison of two groups an unpaired two-tailed Student’s *t*-test or a two-tailed Mann-Whitney *U* test were used. For comparison of two or more factors for each group and their interaction, an ordinary one-way analysis of variance (ANOVA) followed by Tukey’s multiple comparisons test, or a Kruskal-Wallis followed by Dunn’s multiple comparisons test were used. *P* values are indicated as: * *P* < 0.05, ** *P* < 0.005, *** *P* < 0.001, **** *P* < 0.0001.

## Results and Discussion

### CHO cells produce highly abundant TERRA and ARIA transcripts

To characterize the molecular features of telomeric repeat-containing transcripts in CHO cells, we first established a protocol for their depletion using antisense gapmer oligonucleotides (ASOs) complementary to the (UUAGGG)n and (CCCUAA)n sequences, targeting TERRA and ARIA, respectively (α-TERRA and α-ARIA ASOs). Cells were transfected twice, 24 hours apart, with α-TERRA and α-ARIA ASOs, as well as with a non-targeting control (Ctrl) ASO. Total RNA was collected 24 hours after the second transfection and subjected to northern blot hybridization using two strand-specific radiolabeled telomeric probes. Hybridization signals from both probes were extremely strong and appeared as smeary patterns extending from the wells to around 0.5 kb for TERRA, and from above 9 kb to below 0.5 kb for ARIA (Fig. 1A). RNase A treatment confirmed that the detected signals derived from RNA (Fig. 1A). Ctrl ASOs did not alter TERRA or ARIA signals relative to untransfected cells, whereas α-TERRA and α-ARIA ASOs efficiently depleted their respective target RNAs. (Fig. 1A). Given the exceptionally high intensity of the hybridization signals, we conclude that most of these RNA species originate from transcription of the C-rich or the G-rich strand of large heterochromatic ITSs, rather than from telomeres. Whether ITSs are transcribed bidirectionally or whether different ITSs are transcribed in a single, specific direction remains to be determined.

**Figure 1:**
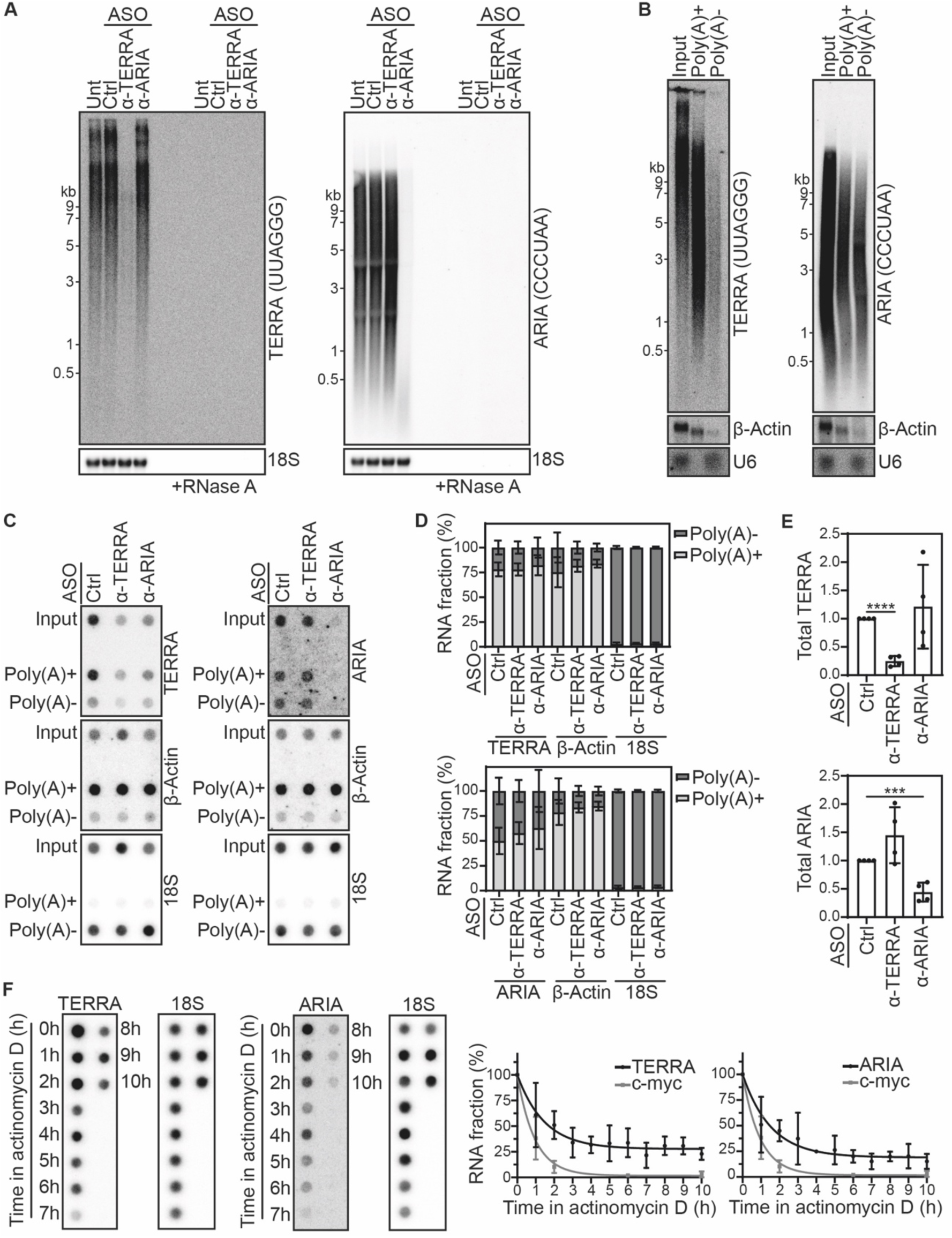
Molecular features of TERRA and ARIA in CHO cells. (**A**) Northern blots detecting TERRA (left) and ARIA (right) using total RNA from cells transfected with ASOs or left untransfected (Unt). Cells were transfected with α-TERRA, α-ARIA and control (Ctrl) ASOs twice, 24 hours apart and RNA was collected 24 hours after the second transfection. Molecular weight markers are shown in kb. Membranes were stripped and re-probed to detect 18S rRNA as a loading control. To confirm signal specificity, RNase A-treated samples were included. (**B**) Northern blots detecting TERRA and ARIA using total (Input), poly(A)+ and poly(A)- RNA fractions. Equivalent RNA amounts were loaded for each fraction. Membranes were stripped and re-probed to detect poly(A)+ β-Actin mRNA and poly(A)- U6 snRNA as controls for loading and fractionation efficiency. (**C**) Dot-blots detecting TERRA and ARIA using total (Input), poly(A)+ and poly(A)- RNA fractions from cells transfected with ASOs as in **A**. Equivalent RNA amounts were loaded for each fraction. Membranes were stripped and re-probed to detect poly(A)+ β-Actin mRNA and poly(A)- 18S rRNA as controls for loading and fractionation efficiency. (**D**) Quantifications of poly(A)+ and poly(A)- RNAs from dot-blot hybridizations as in **C**, expressed as the fraction of the sum of poly(A)+ and poly(A)- signals for each transcript. Bars and error bars are means and standard deviations from four independent experiments. (**E**) Quantifications of total TERRA and ARIA levels using input-associated signals from dot-blot hybridizations as in **C**. Values are normalized to the corresponding 18S signals and expressed as fold increase over ASO Ctrl samples. Bars and error bars are means and standard deviations from four independent experiments. *P* values were calculated using an unpaired two-tailed Student’s *t*-test. (**F**) Dot-blot hybridizations detecting TERRA and ARIA using total RNA from CHO cells treated with actinomycin D for the indicated hours (h). Membranes were stripped and re-probed to detect 18S rRNA as a control for loading. The graphs on the right are quantifications of dot-blots for TERRA and ARIA and of RT-qPCRs for the short-lived c-myc mRNA. TERRA and ARIA signals were normalized to the corresponding 18S signals and GAPDH mRNA was quantified to normalize c-myc. The same c-myc data are shown in the two graphs. Values are normalized to time 0 (set to 100%). Values and error bars are means and standard deviations from four independent experiments for TERRA and ARIA and three independent experiments for c-myc.

We then subjected total RNA to affinity purification using oligo(dT), followed by quantification of TERRA, ARIA, U6 snRNA or 18S rRNA (poly(A)- transcripts) and β-Actin mRNA (poly(A)+ transcript) by northern and dot-blot hybridization. Approximately 75% of TERRA and 50% of ARIA transcripts were detected in the poly(A)+ fraction (Fig. 1B–D). As expected, U6 and 18S RNAs were detected exclusively in the poly(A)- fractions, whereas ∼80% of β-Actin mRNA was recovered in the poly(A)+ fraction, with a residual signal in the poly(A)- fraction (Fig. 1B–D). These results indicate that, although the fractionation protocol was efficient, it was not fully quantitative; therefore, the actual proportion of polyadenylated TERRA and ARIA transcripts might be even higher than estimated. Depletion of TERRA or ARIA did not significantly alter the amounts of the complementary RNA found in either total, poly(A)+ or poly(A)- fractions (Fig. 1C–E). Overall, these results suggest that the majority of TERRA and ARIA transcripts in CHO cells are 3ʹ end-polyadenylated, in contrast to observations in human, mouse and yeast cells, where only a minor fraction of TERRA and ARIA is polyadenylated (21,23,24,36,40). These differences in polyadenylation states likely reflect the predominantly intrachromosomal origin of TERRA and ARIA in CHO cells. Moreover, since depletion of TERRA does not affect the abundance or polyadenylation state of ARIA, and vice versa, the transcription and termination machineries producing these two RNA species likely function independently.

We also measured the half-lives of TERRA and ARIA by treating CHO cells with the global transcription inhibitor actinomycin D and collecting RNA every hour over a 10-hour time course. RNA levels were assessed by dot-blotting for TERRA and ARIA, and by RT-qPCR for the short-lived c-Myc mRNA. While c-Myc levels had already dropped by more than 50% at the first time point, TERRA and ARIA levels declined to approximately 50% after 2 hours of treatment and then stabilized at around 20–30% for the remainder of the time course (Fig. 1F). We speculate that this behavior reflects the existence of two populations of TERRA and ARIA: a more abundant species that is rapidly degraded, with a half-life around 2 hours, and a less abundant but more stable species. Because the majority of CHO TERRA and ARIA transcripts are polyadenylated (Fig. 1B–D), it is unlikely that the polyadenylation state defines the stability of the two populations, as it is the case in human cells (23).

### TERRA and ARIA form large foci in CHO cells

To investigate the cellular localization of TERRA and ARIA, we performed RNA FISH using fluorescently labeled, strand-specific telomeric probes and found that both RNAs were readily detected as large foci, further highlighting their high expression. TERRA foci were more abundant and distributed across both the nucleus and cytoplasm, whereas ARIA foci were approximately half as numerous and nearly exclusively nuclear (Fig. 2A and B). The signals for both TERRA and ARIA were strongly reduced upon ASO transfection and abolished by RNase A treatment, confirming their specificity (Fig. 2A and B). Notably, depletion of TERRA did not visibly affect ARIA foci, while ARIA depletion led to a reduction in the number of TERRA foci (Fig. 2A and B).

**Figure 2:**
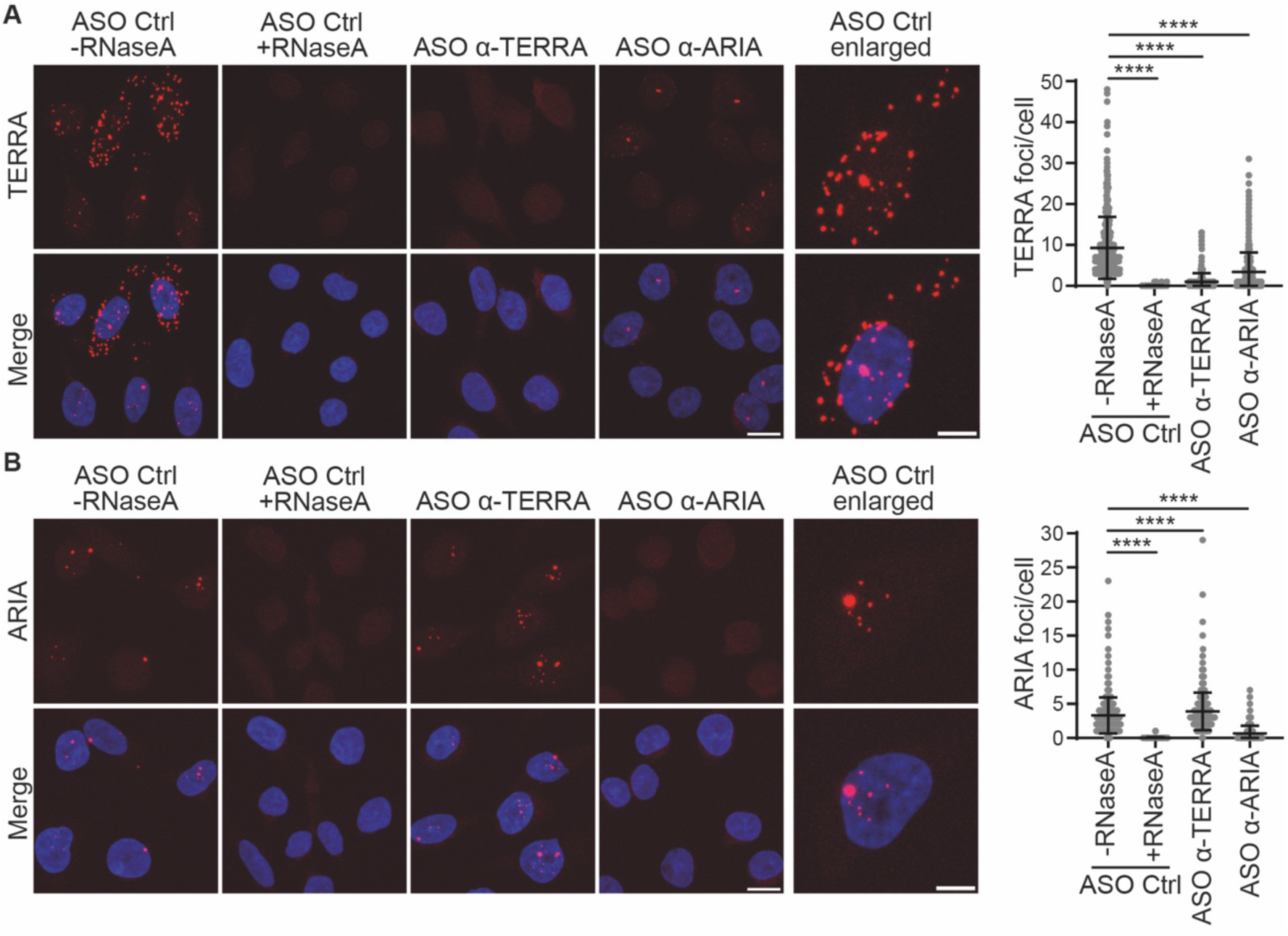
Cellular localization of TERRA and ARIA in CHO cells. Examples of RNA FISH detecting TERRA (**A**) and ARIA (**B**) transcripts in cells transfected with the indicated ASOs. TERRA and ARIA are shown in red, DAPI-stained DNA in blue. To confirm signal specificity, RNase A-treated ASO Ctrl samples were included. Scale bars: 10 µm. Enlarged single nuclei are also shown to facilitate visualization. Scale bars: 5 µm. The plots on the right are quantifications of the numbers of TERRA and ARIA foci per cell from four independent experiments for samples not treated with RNase A and three for RNase A-treated ones. For TERRA FISH, at least 80 nuclei were scored per sample in each experiment for a total of at least 400 nuclei (300 nuclei for RNase A-treated samples). For ARIA FISH, at least 60 nuclei were scored per sample in each experiment for a total of at least 400 nuclei (288 nuclei for RNase A-treated samples). Each dot represents one cell. Means and standard deviations are indicated. *P* values were calculated using a two-tailed Mann-Whitney *U* test.

The nuclear localization of TERRA and ARIA is in agreement with previous reports across multiple organisms for TERRA and with one available cytological analysis in fission yeast for ARIA (18,21,36,41). In addition to that, several observations in mammalian cells, including the association of TERRA transcripts with mitochondria and TERRA non-ATG translation (42,43), indicate that a fraction of TERRA can also localize to the cytoplasm. In CHO cells, where TERRA levels are particularly high, this cytoplasmic pool may be more readily detected. It is also plausible that TERRA transcribed from ITSs is actively transported to the cytoplasm, where it may accumulate in RNA-containing structures such as P-bodies or stress granules (44). Moreover, the reduction in TERRA foci upon ARIA depletion, despite unchanged total TERRA levels (Fig. 1A and E), suggests that ARIA may stabilize a specific fraction of TERRA, likely corresponding to a less soluble pool, as our RNA FISH protocol includes permeabilization and removal of soluble material. We also cannot exclude that this observation may derive from a technical issue, such as interference between the C-rich telomeric probe used to detect TERRA and the G-rich ASO used to deplete ARIA. However, we consider this explanation unlikely, as a similar interference would be expected between the probe detecting ARIA and the α-TERRA ASO.

### ARIA depletion affects ITS stability and cell fitness

To directly test whether TERRA and ARIA telomeric repeat DNA stability, we performed telomeric DNA FISH on metaphase chromosomes and evaluated ITS integrity in ASO-transfected cells. Consistent with previous reports, strong telomeric signals were readily detected at ITSs, whereas telomeric signals at chromosome ends were below the detection limit when applying the same microscopy settings used for ITSs (12,16). Compared with Ctrl and α-TERRA ASO samples, α-ARIA ASOs caused a slight, although not statistically significant, increase in ITS displaying more diffuse and less crisp fluorescence signals (Fig. 3A). This phenotype is reminiscent of fragile telomeres, which are shredded or stretched telomeres found in cells experiencing replication stress (45). We then performed in-gel Southern blot hybridization of genomic DNA digested with a mix of restriction enzymes expected to generate terminal telomeric fragments of 1 kb or less. Gels were hybridized using radiolabeled, strand-specific telomeric repeat probes, first under native conditions and then after denaturation of the same gels to visualize single-stranded (ss) and ds telomeric DNA repeats, respectively. In both conditions, most of the hybridization signals extended from the wells down to 1 kb, indicating that they derived mainly from ITS DNA (11). No major differences across Ctrl, α-TERRA and α-ARIA ASO samples were observed under denaturing conditions (Fig. 3B), suggesting that TERRA and ARIA depletion do not cause gross defects in het-ITS stability. However, ARIA depletion consistently led to a 2–3-fold increase in TTAGGG- and CCCTAA-containing ssDNA (Fig. 3B), a hallmark often associated with processing of DSBs and replication stress (46). Because het-ITS defects, detected by DNA FISH and Southern blotting, occurred specifically upon ARIA but not TERRA depletion, they are unlikely to result from nonspecific interference of telomeric ASOs with DNA replication and instead represent genuine consequences of ARIA insufficiency.

**Figure 3:**
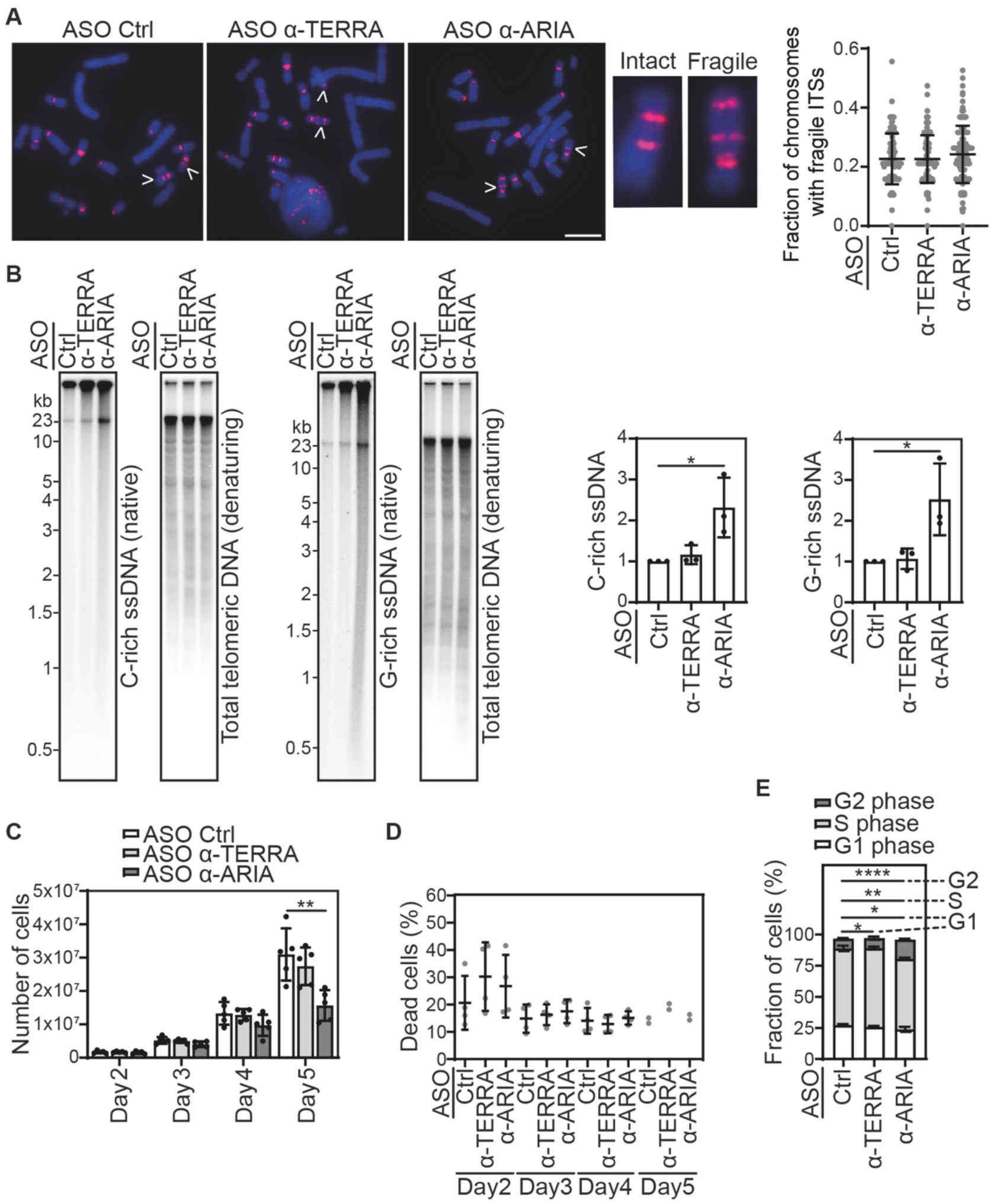
ITS instability and cell proliferation defects in ARIA-depleted CHO cells. (**A**) Examples of telomeric DNA FISH on metaphase chromosomes in cells transfected with the indicated ASOs. Telomeric DNA is shown in red, DAPI-stained DNA in blue. Arrows point to chromosomes with fragile ITSs. Two chromosomes, one with intact and one with fragile ITSs, are enlarged to facilitate visualization. Scale bar: 10 µm. The plot on the right is a quantification of the fraction of chromosomes with fragile ITSs from four independent experiments. At least 21 metaphases were scored per sample in each experiment for a total of at least 100 metaphases. Each dot represents one metaphase. Means and standard deviations are indicated. (**B**) In-gel hybridizations detecting telomeric repeats using total genomic DNA from cells transfected with the indicated ASOs. Gels were first hybridized under native conditions with strand-specific probes to detect single-stranded C-rich and G-rich telomeric repeats. Gels were then denatured and re-probed to detect total telomeric DNA. Molecular weight markers are shown in kb. The graphs on the right are quantifications of ssDNA normalized to the corresponding total DNA and expressed as fold increase over ASO Ctrl samples. Bars and error bars are means and standard deviations from three independent experiments. *P* values were calculated using an unpaired two-tailed Student’s *t*-test. (**C**) Quantifications of total numbers of ASO-transfected cells counted at the indicated times. Bars and error bars are means and standard deviations from five independent experiments. *P* values were calculated using an unpaired two-tailed Student’s *t*-test. (**D**) Quantifications of propidium iodide (PI)-permeable cells (dead cells) in samples transfected with ASOs for the indicated times, expressed as fraction (%) of total cells. Data are from four independent experiments for days 2, 3 and 4 and two for day 5. Each dot represents one experiment. Means and standard deviations are indicated. (**E**) Quantifications of the fraction (%) of ASO-transfected cells in different cell cycle phases based on PI staining of fixed cells. Bars and error bars are means and standard deviations from four independent experiments. *P* values were calculated using an unpaired two-tailed Student’s *t*-test.

Next, we assessed the impact of TERRA and ARIA depletion on cell proliferation and viability. After 5 days of ASO transfections, ARIA-depleted cells were substantially fewer than TERRA-depleted or Ctrl cells (Fig. 3C). Viability assays using propidium iodide (PI) staining of non-permeabilized cells followed by FACS analysis showed no increase in dead cells upon TERRA or ARIA depletion (Figs. 3D and S1A). In contrast, cell cycle analysis of PI-stained, permeabilized cells showed a mild but consistent decrease in S-phase cells and a corresponding increase in G2-phase cells upon ARIA depletion (Figs. 3E and S1B), explaining the lower cell numbers in the same samples. We propose that the accumulation of ARIA-depleted cells in G2 phase is, at least in part, due to activation of a G2/M DNA damage checkpoint triggered by ITS instability. In contrast, TERRA appears to be largely dispensable for ITS integrity maintenance and proliferation at least in unchallenged CHO cells.

### ARIA prevents ssDNA accumulation at damaged ITSs

CHO ITSs are prone to spontaneous breakage because of their repetitive nature (14), leading us to hypothesize that the formation of ssDNA observed upon ARIA depletion might occur at stochastic DSBs within het-ITSs. To test this possibility, CHO cells were treated with camptothecin (CPT), a topoisomerase I inhibitor that generates genome-wide DSBs. Western blot analysis indicated that CPT treatment induced DNA damage, as shown by the accumulation of the DSB marker γ-H2AX, and that ARIA depletion did not alter CPT-induced γ-H2AX levels compared with control cells (Fig. S2A). In native Southern blot analysis, CPT treatment did not detectably affect telomeric ssDNA levels in ASO Ctrl cells. However, combining CPT treatment with ARIA depletion further increased both C-rich and G-rich telomeric ssDNA compared with ARIA depletion alone (Fig. S2B). We next performed native DNA FISH on the same cells using a G-rich telomeric probe to detect C-rich telomeric ssDNA. Consistent with the Southern blot results, few ssDNA foci were observed in Ctrl ASO cells, whereas α-ARIA ASO transfection led to an increase in both the number and intensity of detectable foci. CPT treatment alone did not affect the number or intensity of ssDNA foci compared to untreated samples. In contrast, the combination of CPT treatment and ARIA depletion increased the number and, to a lesser extent, the intensities of detectable foci compared to ARIA depletion alone (Fig. S2C).

To specifically induce DSBs at telomeric repeats, we employed a fusion protein comprising the human shelterin protein TRF1 and the endonuclease FokI (TRF1-FokI), which binds and cleaves TTAGGG dsDNA (47). TRF1-FokI, N-terminally fused to green fluorescent protein (GFP) and a Flag tag, was cloned into a lentiviral vector under the control of a doxycycline (dox)-inducible promoter. CHO cells were infected with lentiviruses encoding TRF1-FokI or control lentiviruses expressing GFP alone, and transgene expression and activity were analyzed after 24 hours of dox treatment. Western blot analysis using an anti-GFP antibody confirmed transgene expression (Fig. S3A), and anti-Flag IF combined with telomeric DNA-FISH showed that TRF1-FokI largely colocalized with ITSs (Fig. S3B). Furthermore, western blotting and double IF using antibodies against γ-H2AX and Flag confirmed that TRF1-FokI induced DNA damage at ITSs (Fig. S3C and D). In native Southern blotting, TRF1-FokI induced the accumulation of both C-rich and G-rich telomeric repeat ssDNA compared with control GFP cells, and ARIA depletion further enhanced ssDNA accumulation in TRF1-FokI-expressing cells relative to either TRF1-FokI or ARIA depletion alone (Fig. 4A). This was even more evident in native DNA FISH, where TRF1-FokI induced the appearance of foci much brighter than the almost undetectable ones observed in GFP control cells (Fig. 4B). The combination of ARIA depletion and TRF1-FokI expression increased both the number and, even more strikingly, the intensity of C-rich ssDNA foci compared with cells expressing TRF1-FokI alone (Fig. 4B). The prominent size and intensity of the foci indicate that the majority of the detected telomeric ssDNA derived from ITSs. Overall, data from CPT- and TRF1-FokI-treated cells support a role for ARIA in regulating damage signaling or repair at damaged telomeric repeat DNA.

**Figure 4:**
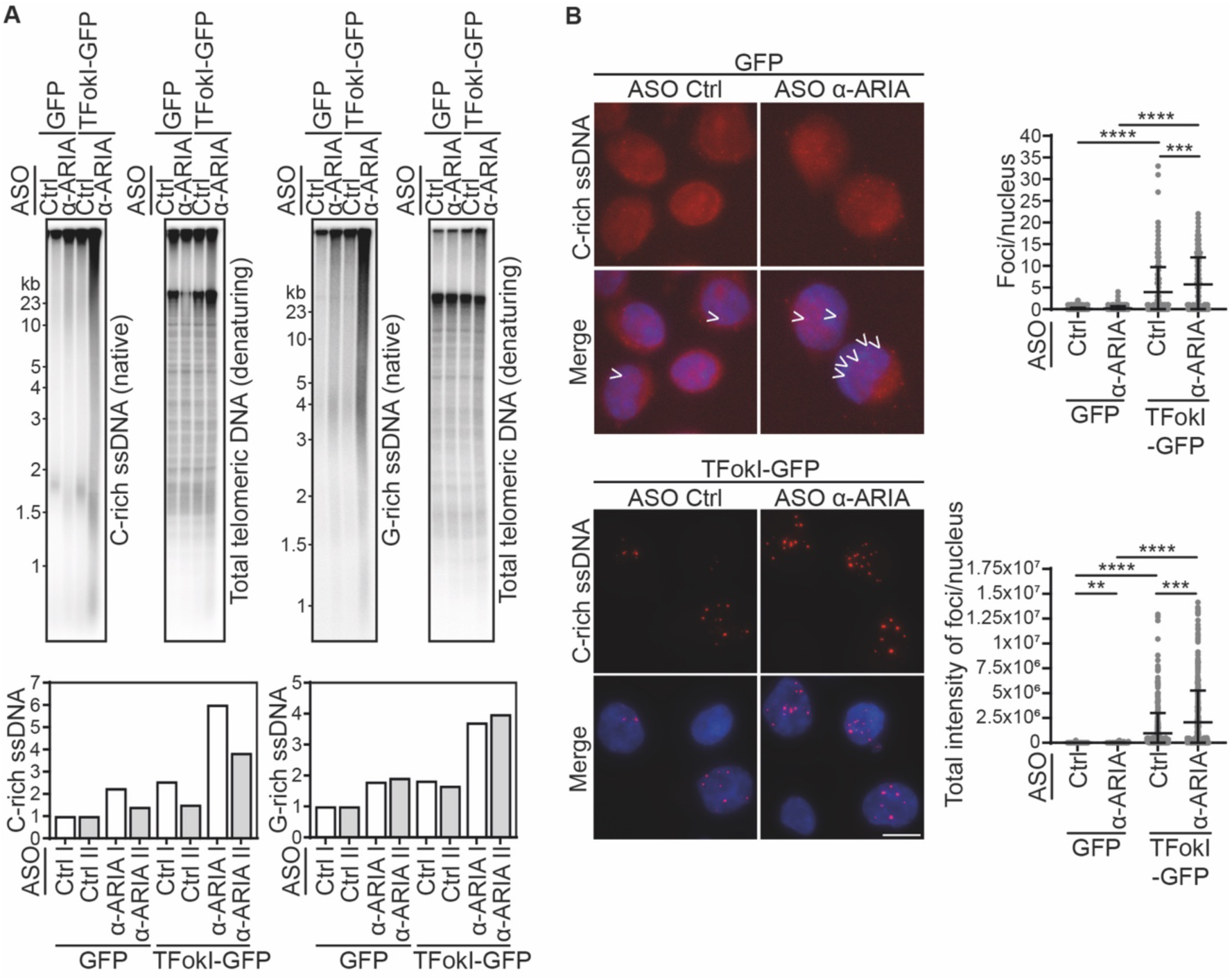
Single-stranded DNA accumulation at damaged ITSs in ARIA-depleted CHO cells. (**A**) In-gel hybridizations detecting single-stranded and total ITS DNA in cells infected with lentiviruses expressing TRF1-FokI-GFP (TFokI-GFP) or GFP only, and transfected with the indicated ASOs. Molecular weight markers are shown in kb. The graphs at the bottom are quantifications of ssDNA normalized to the corresponding total DNA and expressed as fold increase over ASO Ctrl samples from two independent experiments plotted separately (experiments I and II). (**B**) Examples of native DNA FISH detecting C-rich telomeric ssDNA in cells as in **A**. Telomeric DNA is shown in red, DAPI-stained DNA in blue. Images of GFP-infected control cells are displayed at higher brightness than TFokI-GFP images to facilitate visualization of the small ssDNA foci. Scale bar: 10 µm. The plots on the right are quantifications of the numbers and total intensity of telomeric ssDNA foci per nucleus from three independent experiments. At least 68 nuclei were scored per sample in each experiment for a total of at least 300 nuclei. Each dot represents one nucleus. Means and standard deviations are indicated. *P* values were calculated using a Kruskal-Wallis followed by Dunn’s multiple comparisons test.

### ATM, ATR, DNA2 and EXO1 are dispensable for ssDNA generation in ARIA-depleted cells

To unveil the molecular pathways underlying ssDNA accumulation at ITSs in ARIA-depleted cells, we first utilized inhibitors of the DNA damage signaling kinases ATM and ATR (ATMi and ATRi). Western blotting using antibodies against γ-H2AX and phosphorylated CHK1 (p-CHK1) confirmed that both inhibitors efficiently perform in CHO cells. We treated TRF1-FokI-expressing cells transfected with Ctrl or α-ARIA ASOs with ATM or ATR inhibitors (ATMi and ATRi). In native DNA FISH, ATMi had no effect on the number or intensity of ssDNA foci in Ctrl ASO-transfected cells, while in ARIA-depleted cells, it slightly increased the number of foci and markedly enhanced their intensities compared with vehicle-treated controls (Fig. 5A and B). ATRi reduced overall ssDNA accumulation relative to vehicle-treated cells, but did not prevent the increase in ssDNA observed upon ARIA depletion compared with Ctrl ASO-transfected cells. We next examined the contribution of the nucleases DNA2 and EXO1, which are involved in DNA end resection (46). TRF1-FokI-expressing cells were transfected with two independent siRNAs targeting each nuclease (siDNA2-2, siDNA2-3, siEXO1-2, and siEXO1-3) or with non-targeting control siRNAs (siCtrl). After 24 hours, cells were further transfected with Ctrl or α-ARIA ASOs and incubated for an additional 48 hours. At this time point, RT-qPCR indicated an approximate 50% reduction in DNA2 and EXO1 mRNA levels (Fig. S4B). Native DNA FISH showed that depletion of DNA2 or EXO1 did not prevent accumulation of ITS ssDNA in α-ARIA ASO-transfected cells compared with ASO Ctrl-transfected cells (Fig. 6A and B). Our small-molecule inhibitor- and siRNA-based analyses indicate that ATM, ATR, DNA2, and EXO1 are not required for the generation of ssDNA at ITSs when ARIA is depleted. However, ATM appears to limit further accumulation of ssDNA in ARIA-depleted cells, suggesting the existence of a redundant or backup ATM-dependent pathway that prevents excessive ssDNA formation at damaged ITSs specifically when the ARIA activity is impaired.

**Figure 5:**
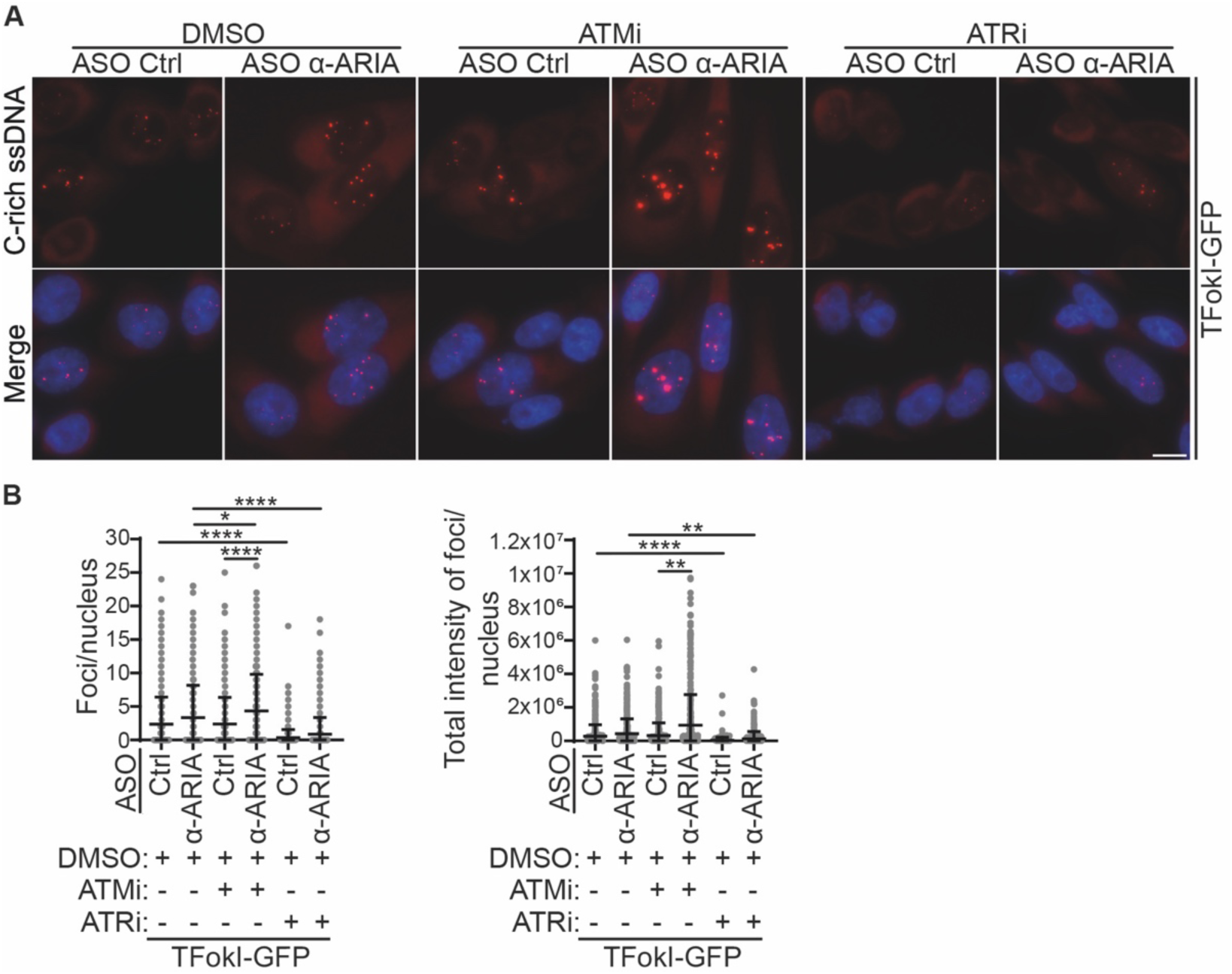
Single-stranded DNA accumulation at damaged ITSs in ARIA-depleted CHO cells treated with ATM and ATR inhibitors. (**A**) Examples of native DNA FISH detecting C-rich telomeric ssDNA in cells expressing TRF1-FokI-GFP (TFokI-GFP) transfected with the indicated ASOs and treated with an ATM inhibitor (ATMi), an ATR inhibitor (ATRi) or with DMSO (vehicle) for 5 hours before harvesting. Telomeric DNA is shown in red, DAPI-stained DNA in blue. Scale bar: 10 µm. (**B**) Quantifications of the numbers and total intensity of telomeric ssDNA foci per nucleus from three independent experiments. At least 80 nuclei were scored per sample in each experiment for a total of at least 300 nuclei. Each dot represents one nucleus. Means and standard deviations are indicated. *P* values were calculated using a Kruskal-Wallis followed by Dunn’s multiple comparisons test.

**Figure 6:**
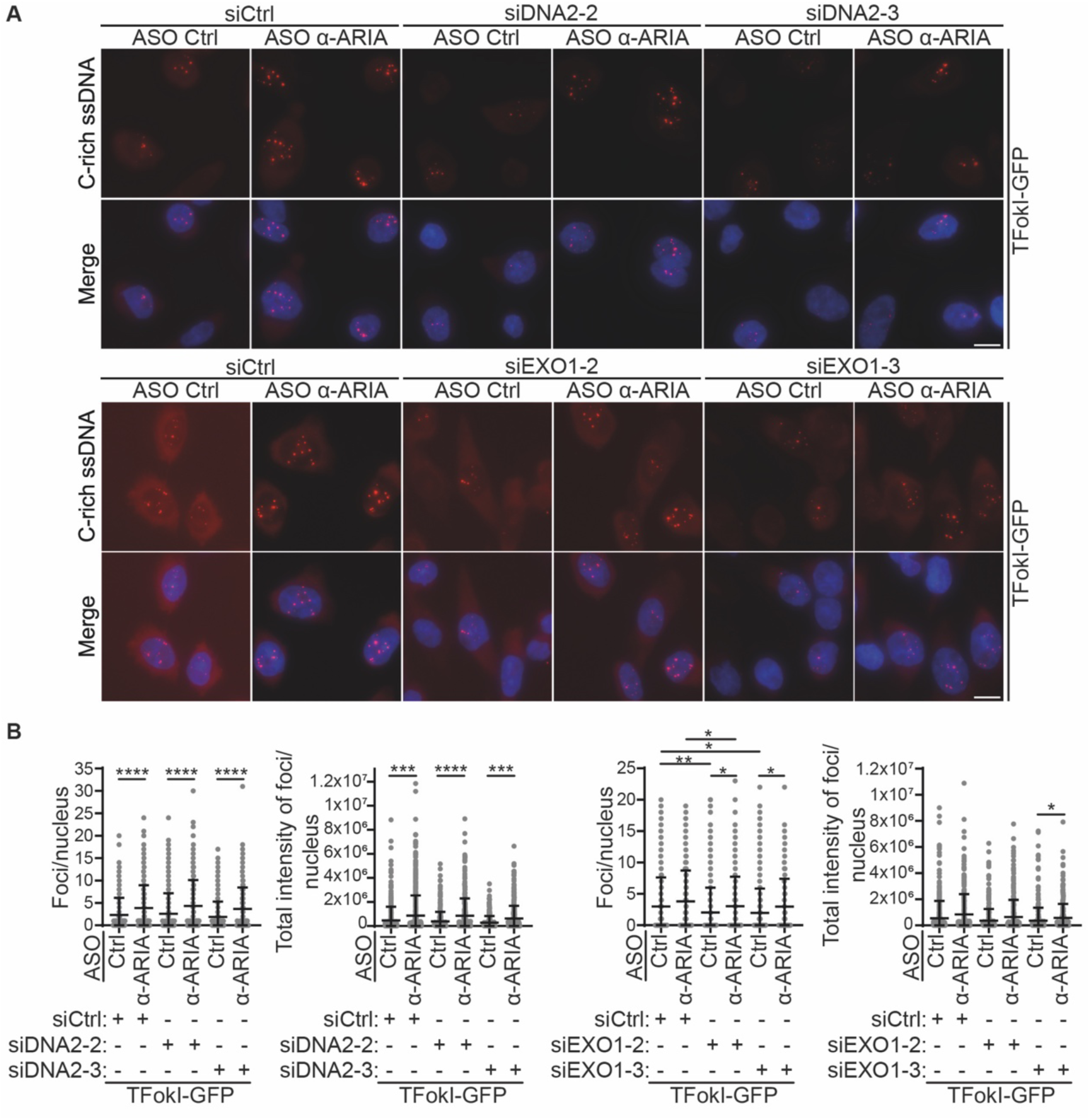
Single-stranded DNA accumulation at damaged ITSs in ARIA-depleted CHO cells depleted of DNA2 and EXO1. (**A**) Examples of native DNA FISH detecting C-rich telomeric ssDNA in cells expressing TRF1-FokI-GFP (TFokI-GFP) transfected with the indicated ASOs and depleted of DNA2 or EXO1 using siRNAs. Cells were first transfected with two independent siRNAs against each factor (siRNAs 2 and 3) or a non-targeting siRNA control (siCtrl). 24 hours later, cells were transfected twice with ASOs 24 hours apart and harvested 24 hours after the second transfection. Telomeric DNA is shown in red, DAPI-stained DNA in blue. Scale bars: 10 µm. (**B**) Quantifications of the numbers and total intensity of telomeric ssDNA foci per nucleus from three independent experiments for DNA2 depletion and two independent experiments for EXO1 depletion. For DNA2 depletion, at least 94 nuclei were scored per sample in each experiment for a total of at least 300 nuclei. For EXO1 depletion, at least 100 nuclei were scored per sample in each experiment for a total of at least 200 nuclei. Each dot represents one nucleus. Means and standard deviations are indicated. *P* values were calculated using a Kruskal-Wallis followed by Dunn’s multiple comparisons test.

### Conclusions

Here, we present the first extensive characterization of telomeric-repeat-containing RNAs transcribed from intrachromosomal telomeric DNA in a mammalian model system. ITSs are also transcribed in other mammals, including humans (22,39); however, the limited length of the ITS tracts and their relatively low expression levels make analysis in those systems more challenging. Our work clearly establishes that CHO cells provide a uniquely amenable system for studying features and functions of telomeric repeat-containing RNAs. Several functions exerted by telomeric RNAs produced from ITSs are expected to be conserved with TERRA and ARIA species produced from telomeres, for example those that depend on interactions with RNA-binding proteins recognizing UUAGGG or CCCUAA repeats. The high abundance of TERRA and ARIA in CHO cells should facilitate the identification of such proteins and the study of their relevance at both ITSs and telomeres across mammalian evolution.

Confirming the utility of CHO cells, our current work uncovers a novel function of ARIA in preventing excessive ssDNA accumulation at damaged telomeric repeat DNA. One conceivable explanation for this mechanism is that ARIA restricts DNA resection at breaks through a pathway that does not involve ATR, DNA2 or EXO1, and which is backed up by ATM. But how does ARIA achieve this? Because TERRA depletion does not cause ssDNA exposure, we exclude the involvement of small RNAs derived from double-stranded dilncRNAs containing G-and C-rich telomeric repeats (38). One possibility is that ARIA forms RNA:DNA hybrids at the break site, thus creating a physical barrier against resection. Alternatively, ARIA could recruit protein inhibitors of resection activities to the breaks. The first step towards understanding the precise mechanism will be to clarify the molecular structure of the ssDNA and identify the enzymatic activities responsible for its generation. It will also be crucial to determine whether this ARIA-associated function is limited to DSBs at ITSs or it also extends to telomeres. Given that the telomeric G-overhang resembles the resected end of a telomeric DSB, ARIA, and perhaps ATM when ARIA-dependent functions are compromised, may play an unappreciated role in its formation. The contribution of ARIA to G-overhang regulation might have been overlooked so far simply because of the very low abundance of this telomeric RNA.

## Data Availability

All data relevant to the study are included in this article or uploaded as supplemental information. If needed, additional data can be requested to the corresponding author.

## Supplementary Data Statement

Supplementary Data are available for this article.

## Acknowledgements

We thank Elena Giulotto for providing CHO cells, Roger Greenberg for providing a plasmid containing TRF1-FokI cDNA, Emilio Cusanelli for sharing the ImageJ macro used for image quantifications, Marta Andolfato for assistance with macro implementation and Patrícia Abreu for critical reading of the manuscript. We also thank the Bioimaging and Flow Cytometry Platforms of GIMM for technical support and the Azzalin laboratory for fruitful discussions.

## Author contributions

B.D-S. and C.M.A. conceptualized the study and wrote the manuscript; B.D-S. performed all experiments and analyzed the data; C.M.A. provided funding.

## Supplementary Figures

**Supplementary Figure S1:**
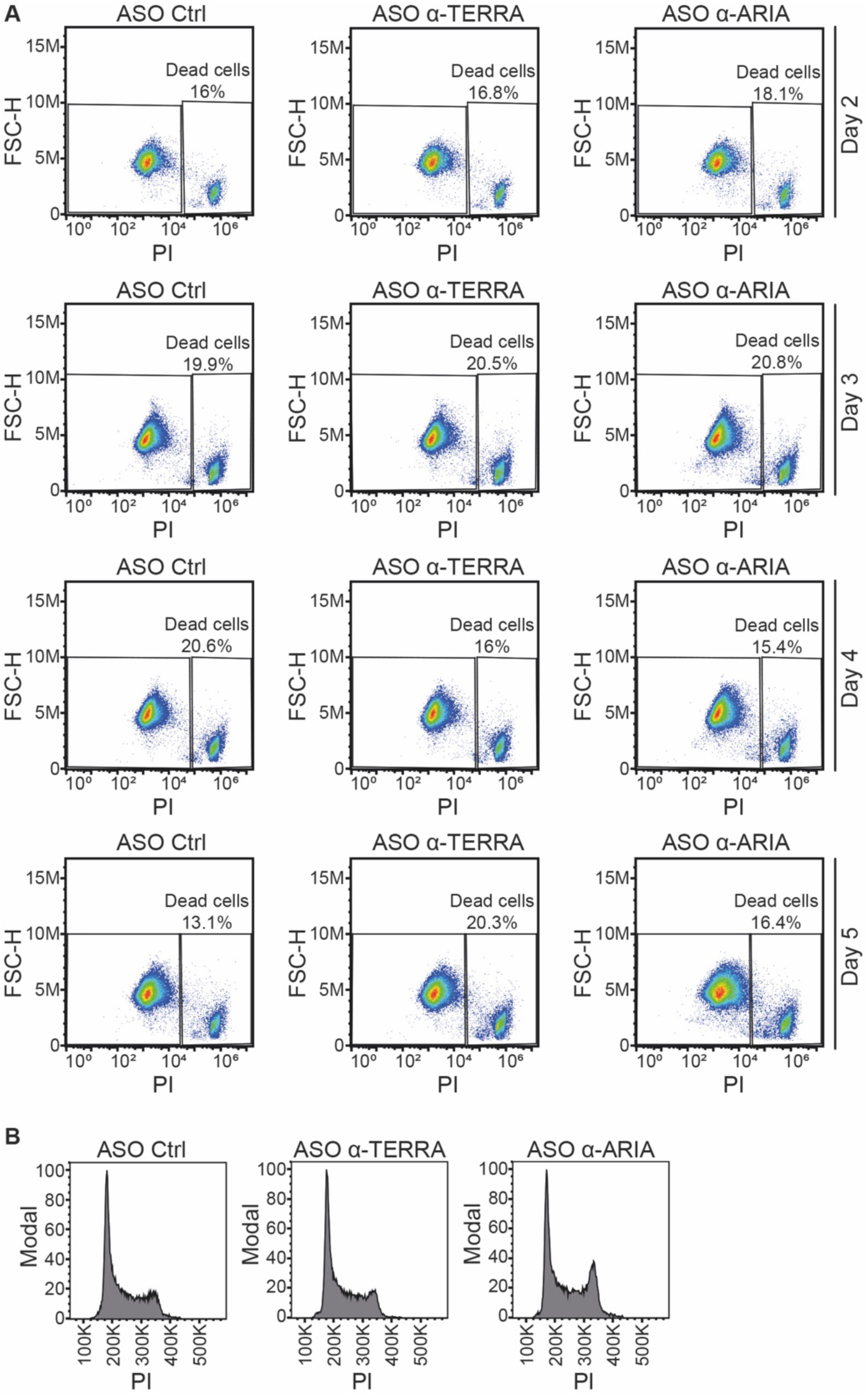
Cell death and cell cycle analysis of TERRA- and ARIA-depleted CHO cells. (**A**) Representative fluorescence-activated cell sorting (FACS) analysis of ASO-transfected cells stained with propidium iodide (PI) in the absence of permeabilization and fixation. The fraction (%) of PI-positive, dead cells is indicated. FSC-H: Forward Scatter-Height. (**B**) Representative FACS analysis of ASO-transfected cells stained with PI after permeabilization and fixation using ethanol.

**Supplementary Figure S2:**
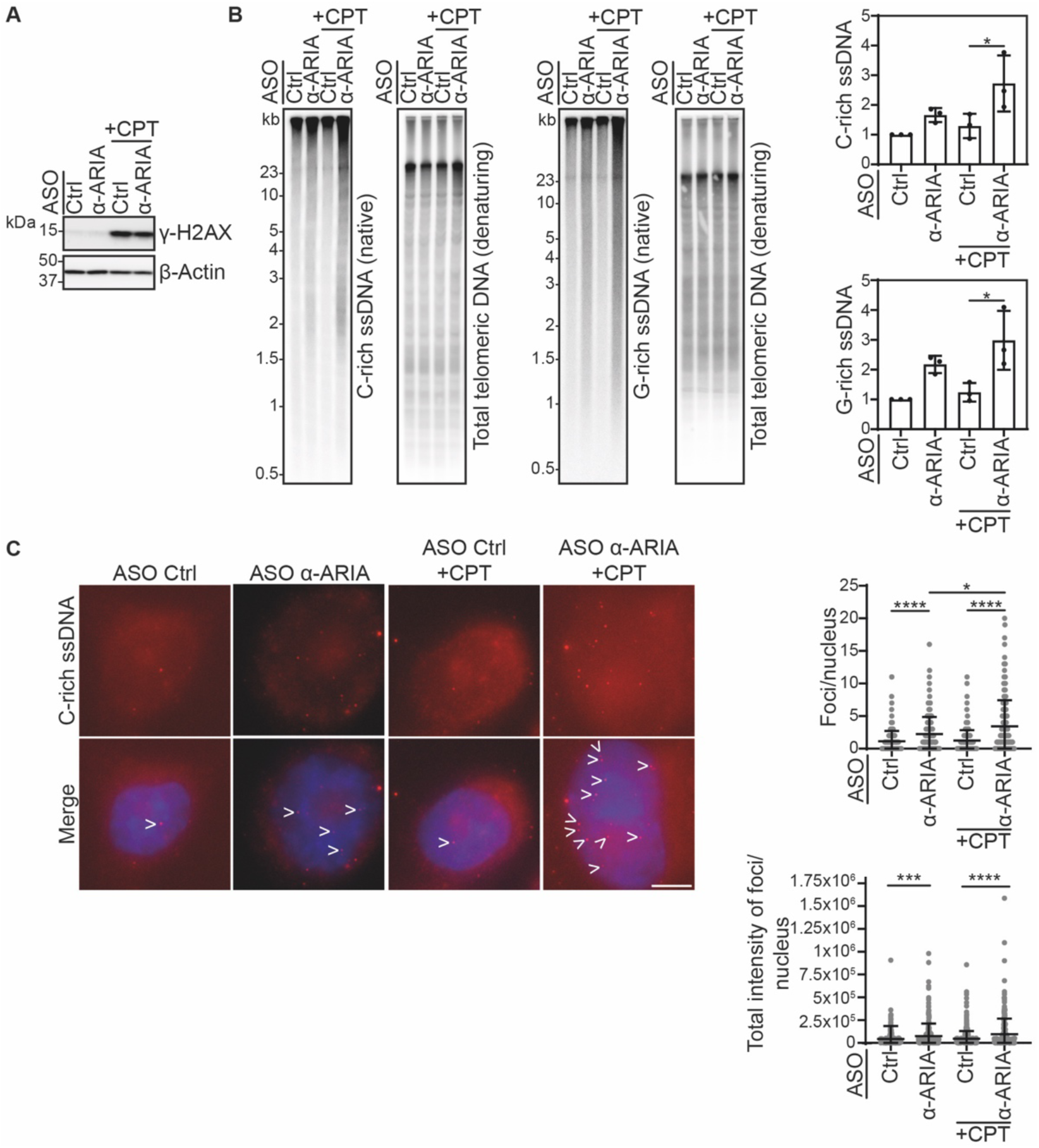
Single-stranded DNA accumulation at ITSs in ARIA-depleted CHO cells treated with camptothecin. (**A**) Western blot detecting γ-H2AX and β-Actin (loading control) using total protein extracts from cells transfected with the indicated ASOs and treated with 150 nM camptothecin (CPT) for 1 hour. Marker molecular weights are shown in kDa. (**B**) In-gel hybridizations detecting telomeric repeats using total genomic DNA from cells as in **A**. Gels were first hybridized under native conditions with strand-specific probes to detect single-stranded C-rich and G-rich telomeric repeats. Gels were then denatured and re-probed to detect total telomeric DNA. Molecular weight markers are shown in kb. The graphs on the right are quantifications of ssDNA normalized to the corresponding total DNA and expressed as fold increase over ASO Ctrl samples. Bars and error bars are means and standard deviations from three independent experiments. *P* values were calculated using an ordinary one-way ANOVA followed by Tukey’s multiple comparisons test. (**C**) Examples of native DNA FISH detecting C-rich telomeric ssDNA in cells as in **A**. Telomeric DNA is shown in red, DAPI-stained DNA in blue. Arrows point to C-rich telomeric ssDNA. Scale bar: 5 µm. The graphs on the right are quantifications of the numbers and total intensity of telomeric ssDNA foci per nucleus from three independent experiments. At least 63 nuclei were scored per sample in each experiment for a total of at least 249 nuclei. Each dot represents one nucleus. Means and standard deviations are indicated. *P* values were calculated using a Kruskal-Wallis followed by Dunn’s multiple comparisons test.

**Supplementary Figure S3:**
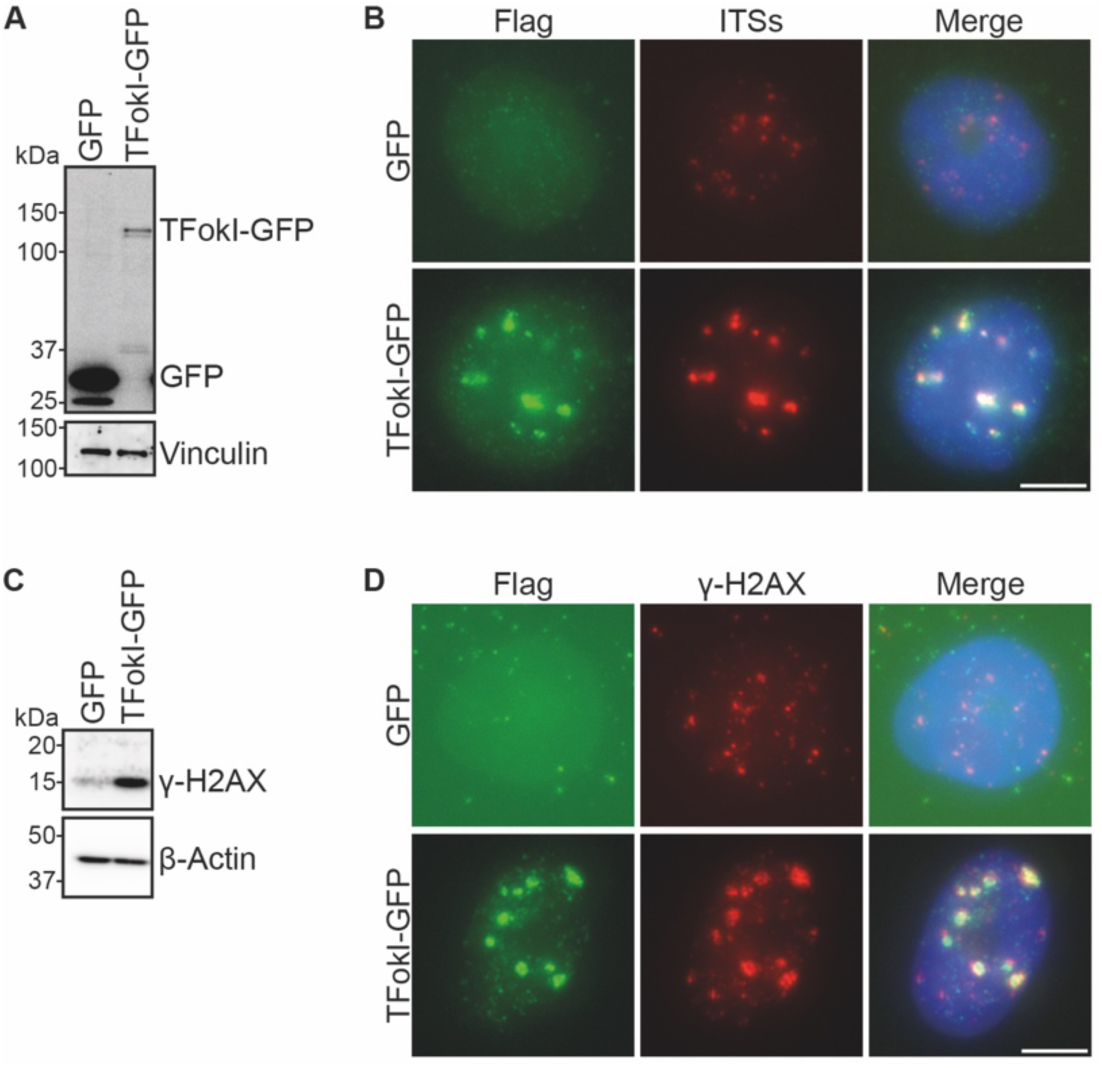
Validation of the TRF1-FokI system in CHO cells. (**A**) Western blot detecting GFP using total protein extracts from cells infected with lentiviruses expressing TRF1-FokI-GFP (TFokI-GFP) or GFP only. The membrane was stripped and probed to detect Vinculin as a loading control. Marker molecular weights are shown in kDa. (**B**) Examples of IF/DNA FISH detecting telomeric DNA and the Flag epitope in cells as in **A**. Flag staining is shown in green, telomeric DNA (ITSs) in red, DAPI-stained DNA in blue. Scale bar: 5 µm. (**C**) Western blot detecting γ-H2AX and β-Actin (loading control) using total protein extracts from cells as in **A**. Marker molecular weights are shown in kDa. (**D**) Examples of double IF detecting the Flag epitope and γ-H2AX. Flag staining is shown in green, γ-H2AX in red, DAPI-stained DNA in blue. Scale bar: 5 µm.

**Supplementary Figure S4:**
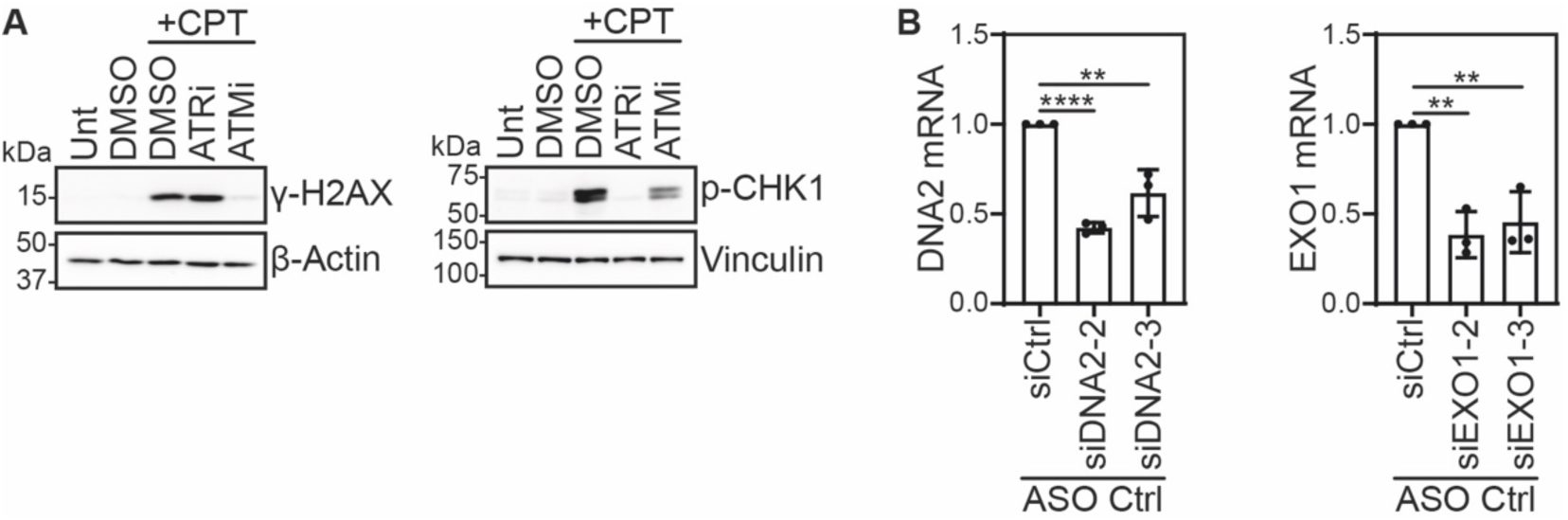
Validation of ATM and ATR inhibitors and of DNA2 and EXO1 siRNAs. **(A)** Western blots detecting γ-H2AX and phosphorylated CHK1 (p-CHK1) using total protein extracts from cells treated with 1 µM CPT for 1 hour on top of treatment with DMSO (vehicle), an ATR inhibitor (ATRi) or an ATM inhibitor (ATMi) for 5 hours before harvesting. β-Actin and Vinculin were used as loading controls. Marker molecular weights are shown in kDa. **(B)** Quantifications of RT-qPCRs for DNA2 and EXO1 mRNAs in cells transfected with two independent siRNAs against DNA2 (siDNA2-2 and siDNA2-3), two independent siRNAs against EXO1 (siEXO1-2 and siEXO1-3) or a non-targeting siRNA control (siCtrl). Cells were harvested 72 hours after transfection. GAPDH mRNA was quantified to normalize DNA2 and EXO1. Values are plotted as fold changes over siCtrl samples. Bars and error bars are means and standard deviations from three independent experiments. *P* values were calculated using an unpaired two-tailed Student’s *t*-test.

